# Bimodal distribution of *Candida albicans* in children with Autism linked with ASD symptoms

**DOI:** 10.1101/2023.07.13.548908

**Authors:** Khemlal Nirmalkar, Jigar Patel, Dae-Wook Kang, Andrew Bellinghiere, Devin A. Bowes, James B. Adams, Rosa Krajmalnik-Brown

## Abstract

The gastrointestinal (GI) tract harbors an intricate and remarkably diverse microbial ecosystem that profoundly impacts various aspects of health and pathophysiology. While bacteria overwhelmingly represent most of the GI microbiota, it is imperative to consider the presence and function of fungal constituents (i.e., mycobiota) within the GI ecosystem. The substantial incidence of GI disorders and associated manifestations in children diagnosed with autism spectrum disorder (ASD) suggests a plausible contributory role of the gut mycobiota. Our investigation aimed to elucidate the gut mycobiota in a cohort comprising 38 typically developing children (TD) and 40 children with ASD. Fecal samples were collected from all participants and autism severity and GI symptoms were assessed to unravel the potential implications of mycobiota alterations in the gut. We employed fungal internal transcribed spacer (ITS) gene amplicon sequencing to analyze the fungal composition and investigate their relationship with GI and autism symptoms. Among gut mycobiota, *Saccharomyces cerevisiae* was significantly lower (relative abundance) in ASD compared to TD children. Total *Candida* and *C. albicans* demonstrated a bimodal distribution among children with ASD. Children with ASD with elevated Autism Treatment Evaluation Checklist (ATEC) scores (a more severe diagnosis) displayed an increased abundance of *C. albicans* and a decreased abundance of *S. cerevisiae*. A significant positive correlation was observed between ATEC scores and GI symptoms and between ATEC scores and *C. albicans*. Our findings propose that a deficit of beneficial fungi, specifically *S. cerevisiae,* and an overgrowth of *C. albicans* may worsen autism severity in children with ASD. Future work employing more advanced techniques (i.e., shotgun metagenomics) is encouraged to advance understanding of the functional role of fungi/yeast, and their interplay between GI symptoms and autism severity in children with ASD.

## Introduction

The incidence of autism spectrum disorder (ASD) within the United States has been rising steadily increasing, and with recent reports by, the Centers for Disease Control and Prevention (CDC), noting 1 in 36 children are diagnosed with ASD ^1^. ASD is a multifactorial neurodevelopmental disorder ^2^ and multiple reports have shown that ASD is associated with gut symptoms ^3^ and abnormal gut microbiota ^4,5^. The gut microbiota affects communication between the gut and brain, which has been found to impact brain function and behavior in ASD ^6,7^.

It has been demonstrated that an imbalance in the gut bacterial composition is present in children with ASD and is linked with ASD and gastrointestinal (GI) symptoms ^4^. Common beneficial bacteria, such as *Bifidobacterium, and Prevotella,* have been found in lower abundance ^5,8–10^, while an increased abundance of *Clostridia, Sutterella, and Bacteroides* have been observed in children with ASD compared to typically developing controls ^11–14^.

Abnormal gut bacteria could result from increased oral antibiotic usage, a common occurrence in children with ASD ^4,9,15^. This combined scenario of excessive antibiotic usage and abnormal gut bacteria could lead to the overgrowth of opportunistic fungi or yeast in ASD ^16^. An increased abundance of fungi or yeast (primarily *Candida*) has been reported in 25-58% of children with ASD, at rates substantially higher than in TD individuals ^3,17–21^. Among fungi, a higher abundance of *Aspergillus* was reported in one ASD child (a case report) ^22^, but in contrast, a lower abundance of *Aspergillus versicolor* was found in 29 ASD children compared to 31 TD ^20^. Whereas, for yeast, a higher abundance of *Candida* specifically *C. albicans* was consistently observed in children with ASD compared to controls ^18,19,21^.

Yeast has been associated with more severe ASD symptoms, presumably due to the production of mycotoxins, such as alcohols and aldehyde ^23–26^, known to adversely impact mental function ^27^ potentially exacerbating existing ASD-related symptoms ^28^. The increase in yeast in ASD children has been found to be associated with lower secretory Immunoglobulin A (sIgA) ^29^, which is the primary antibody combating bacteria and fungi in the mucous membranes ^30,31^. A small open-label clinical trial revealed that treating yeast infections with antifungals led to some clinical improvements in symptoms ^28^, and a national survey of ASD families found that anti-fungal treatments (Nystatin and Diflucan) were rated among the most effective treatments for ASD symptoms ^32^. The high incidence of yeast infections in children with ASD suggests that many children with ASD have long-lasting yeast infections. Given the substantial proportion of children being diagnosed on the spectrum, it is imperative to explore the potential correlation between fungal microbiota and ASD in children.

Hence, this pilot study investigated, gut mycobiota in children with ASD and TD, using fungal ITS sequencing and qPCR targeting the fungal 18S rRNA gene. We looked at differences in key fungal and yeast composition, and analyzed their relationship with GI and ASD symptoms.

## Results

### S. cerevisiae and C. albicans were dominant gut mycobiota

After ITS fungal gene amplicon sequencing of fecal DNA from 38 TD and 40 ASD children (Table S1), we received 5,089,812 reads, averaging 65,254 reads per sample. Qiime2 ^33^ and downstream analyses (refer to methods) did not exhibit any significant difference in alpha (Fig. S1) and beta diversity (Fig. S2) between TD and ASD children. Utilizing the UNITE database ^34^ for taxonomic classification, we identified 146 fungal taxa such as *S. cerevisiae, C. albicans, Penicillium, Candida sake, Cladosporium etc.* (Table S2). Relative abundance calculation (Fig. 1A, Fig.S3) showed that *S. cerevisiae* predominated in the gut mycobiota of TD, whereas *C. albicans* was prominent in ASD group cohort (Fig. 1B), whereas the biplot (Fig. 1B), which indicates overall dominance of microbial features in all samples, and in this study, we observed *S. cerevisiae* and *C. albicans* were the most dominant fungi. Notably, *S. cerevisiae* was significantly lower (p=0.011) in ASD participants as opposed to TD group (Fig. 2A). Conversely, *C. albicans* did not exhibit a statistically significant difference (p>0.05) between ASD and TD (Fig. 2C). However, a fraction, constituting 27.5% (n=11/40) of ASD children, had more than 10% relative abundance of total *Candida* and *C. albicans*.

**Figure 1:**
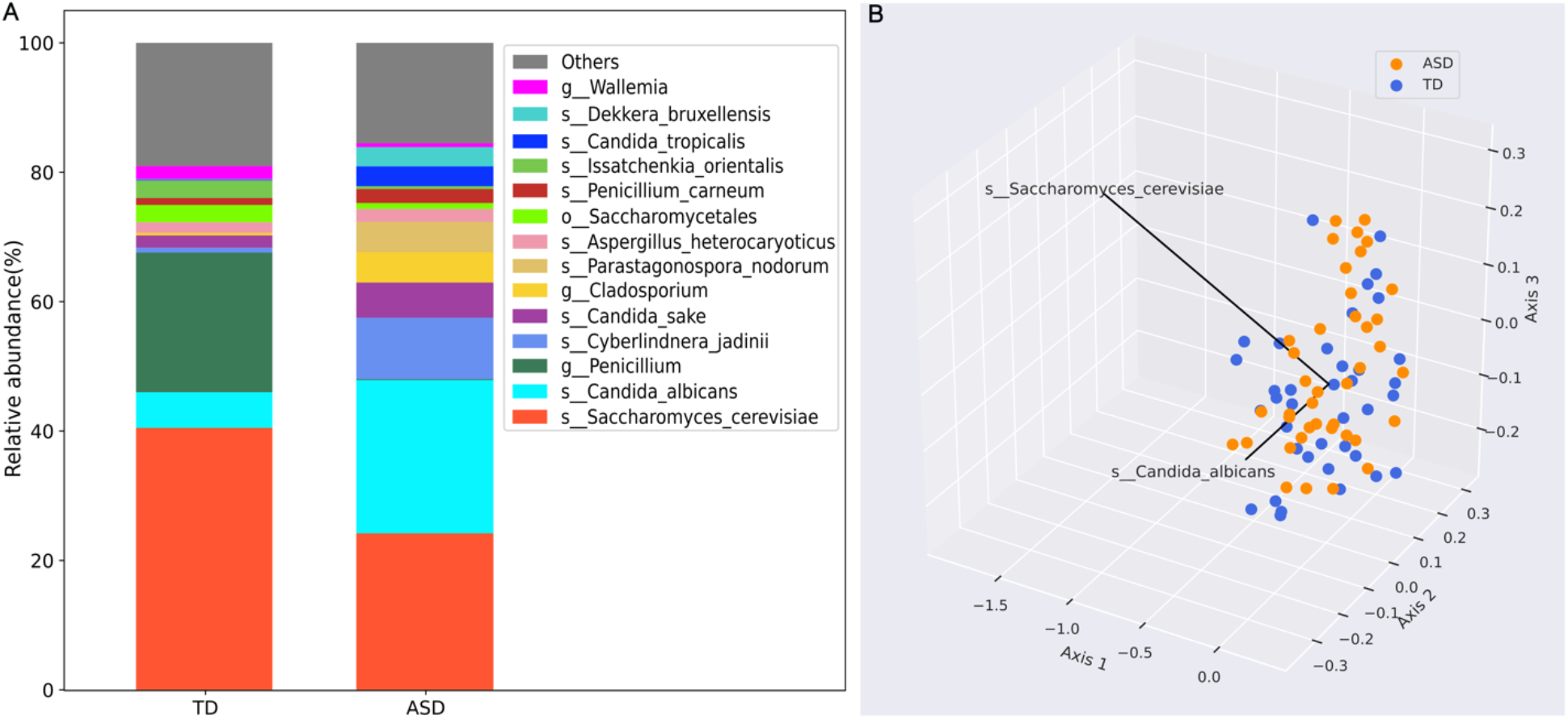
Average relative abundance of Top 15 fungal taxa in TD and ASD children (A) and most dominant fungal microbiota in TD and ASD children using biplot (B). TD-typically developing; ASD-autism spectrum disorder. Blue dots represent TD and orange color for ASD participants. Relative abundance of fungi in each participant, refer to supplementary Figure S4.

**Figure 2:**
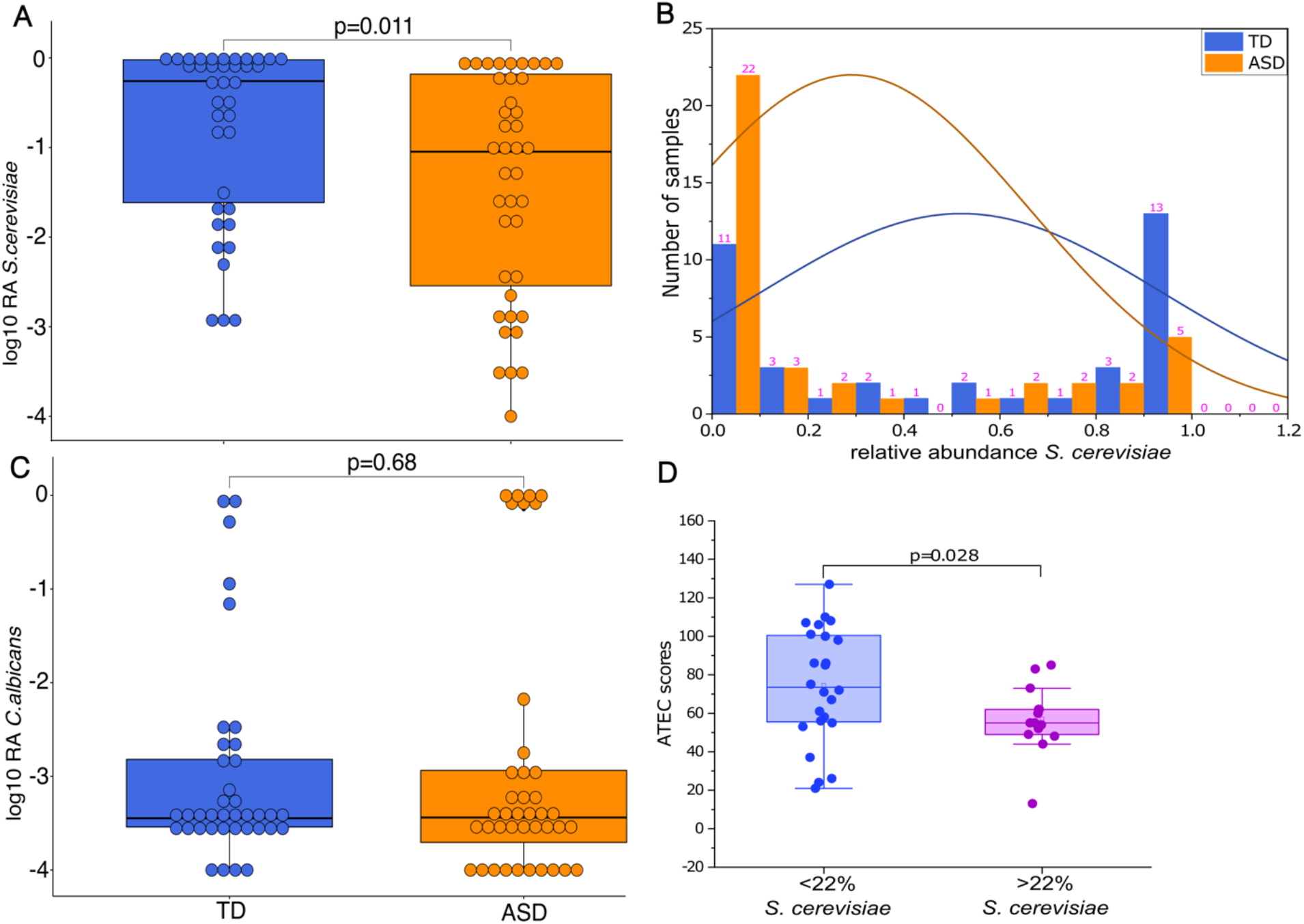
Univariate comparison of the relative abundance (after log10 transformation) of *S. cerevisiae* (A) and *C. albicans* (C). Distribution of *S. cerevisiae* relative abundance in TD and ASD participants (B), ATEC scores comparison between <22% and >22% relative abundance of *S. cerevisiae* in children with ASD (D). TD-typically developing; ASD-autism spectrum disorder. p values < 0.05 are considered statistically significant and taxa p-values are adjusted with the BH correction method.

### Children with ASD presented a bimodal distribution of S. cerevisiae and C. albicans

The distribution of *S. cerevisiae’s* relative abundance (Fig. 2B) revealed a bimodal distribution in ASD children. We used 22% as a cut-off value for the relative abundance of *S. cerevisiae* due to the abundance pattern in ASD children (Fig. 2B). Interestingly, 62.5% of children with ASD exhibited a diminished abundance of S. *cerevisiae* (<22%) and concurrently manifested significantly elevated Autism Treatment Evaluation Checklist (ATEC) (p=0.028) scores (high ATEC scores are a result of a more sever diagnosis) in comparison to those with a higher abundance of *S. cerevisiae* (>22%) (Fig. 2D). Interestingly, in those ASD individuals who had >22% *S. cerevisiae*, their *C. albicans* abundance was <0.01%. The relative abundance of all *Candida* (Fig 3A) and of *C. albicans* (Fig 3B) also showed a bimodal distribution in ASD children. We chose the criteria of 80% relative abundance to define those with elevated relative abundance. The subset of ASD children with a high relative abundance of total *Candida* (>80%), and in that subset (85.7%) of ASD children with a high relative abundance of *C. albicans* (>80%) had significantly higher ATEC scores than their counterparts (p=0.034 *Candida*; p=0.007 *C. albicans*) (Fig. 3C-D). Additionally, to quantify the absolute load of fungi, we performed qPCR targeting the fungal 18S rRNA gene. However, we did not observe a significant difference between TD and ASD (Fig. S4A). Furthermore, the semi-quantitively gene abundance calculation (see methods) of *S. cerevisiae* and *C. albicans* showed no significant difference between TD and ASD children (Fig. S4B-C).

**Figure 3:**
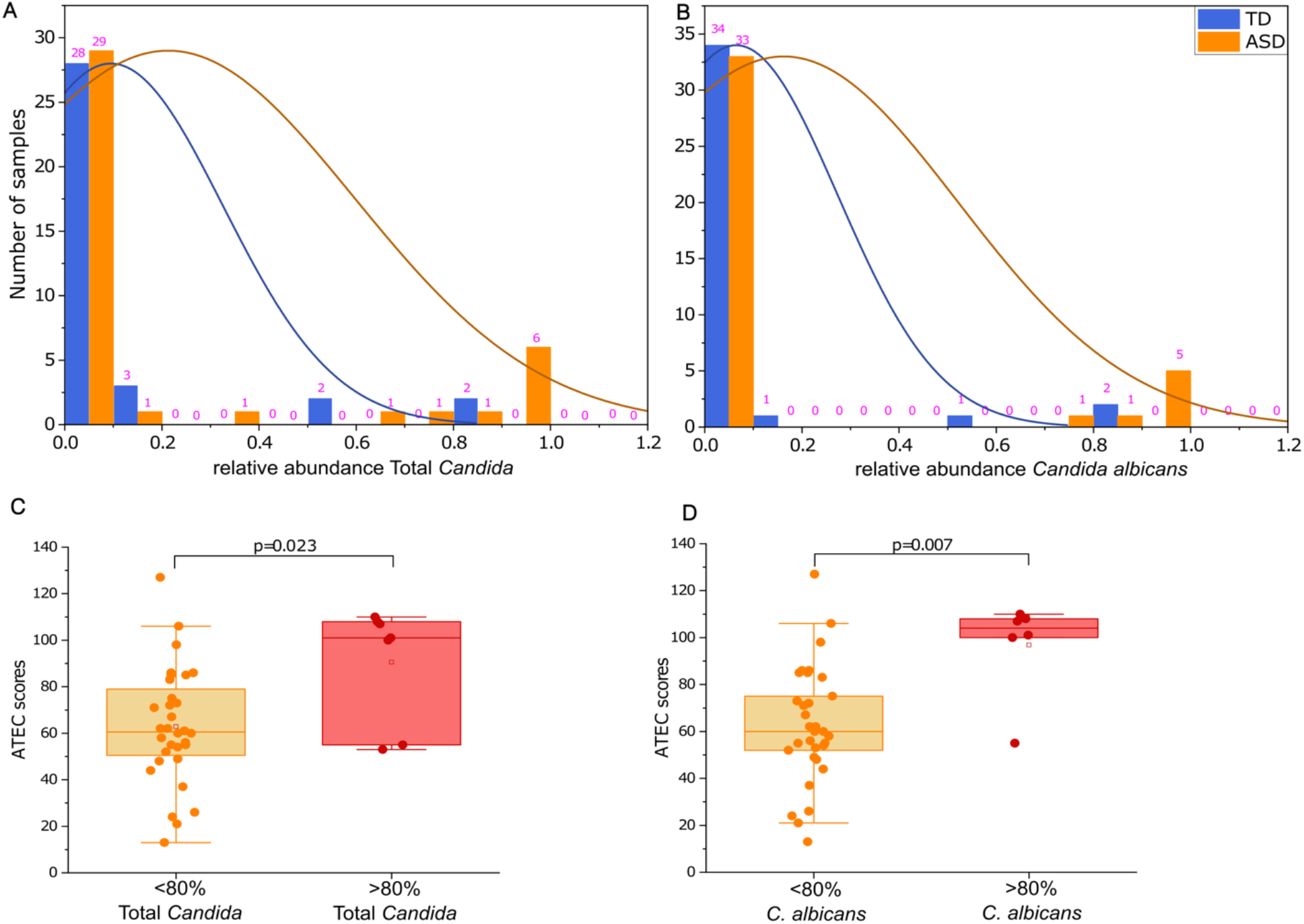
Distribution of total *Candida* (A) and *C. albicans* (B) relative abundance in TD and ASD participants. ATEC scores comparison between <80% and >80% relative abundance of Total *Candida* (C) and *C. albicans* (D) in children with ASD. TD-typically developing; ASD-autism spectrum disorder. p values < 0.05 are considered statistically significant.

### ASD Symptoms are correlated with GI symptoms and C. albicans

We evaluated GI symptoms with six sub-items of the 6-GSI: constipation, diarrhea, consistency, smell, flatulence, and abdominal pain in TD and ASD children. We observed that children in the ASD cohort exhibited significantly higher (p<0.05) GI symptoms in comparison to their TD counterparts (Fig. S5). Comparing ATEC scores for children with ASD with associated GI symptoms, revealed a significantly positive correlation (Spearman R=0.37, p=0.02; Pearson R=0.36, p=0.02) (Fig. 4A). However, ATEC scores did not demonstrate a significant correlation with *S. cerevisiae* (Fig. 4B). Contrastingly, a small but positive correlation between ATEC scores and *C. albicans* was observed (Spearman R=0.23, p=0.15; Pearson R=0.35, p=0.02) (Fig. 4C). Intriguingly, there was a significant negative correlation between *S. cerevisiae and C. albicans* (Spearman R=-0.41, p=0.008; Pearson R=-0.35, p=0.025) (Fig. 4D) in the ASD cohort. Though these correlations are statistically significant, it is important to note that *S. cerevisiae* and *C. albicans* showed a bimodal distribution for ATEC scores (values are in two extreme points). Our analysis did not reveal significant correlations between GI symptoms and either *S. cerevisiae* or *C. albicans* in ASD (Fig. S6A-B), or TD children (Fig. S6C-D). Similarly, *S. cerevisiae* did not significantly correlate with *C. albicans* in the TD cohort.

**Figure 4:**
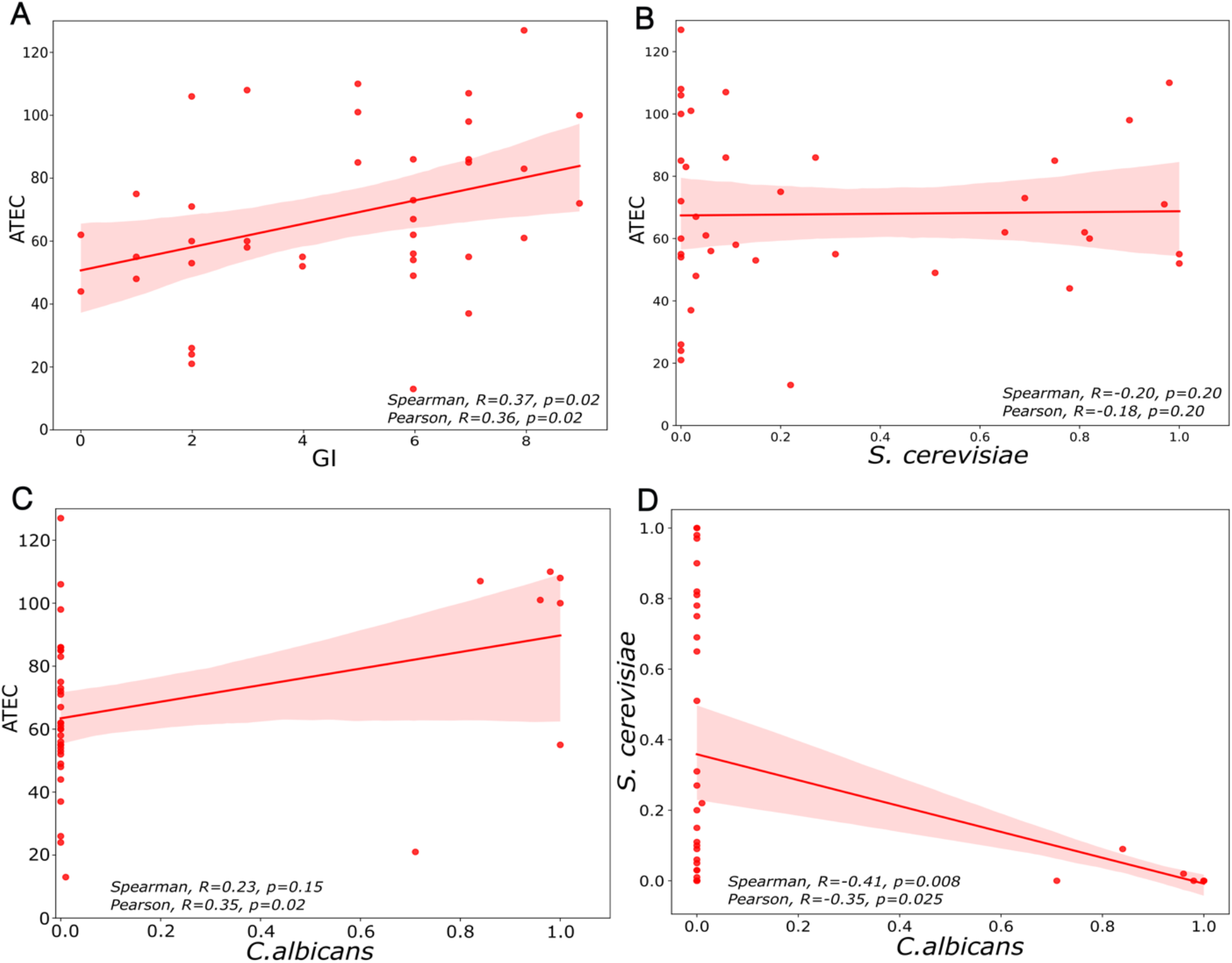
Correlation analyses between fungal taxa and symptoms in children with ASD. Between ATEC and GI scores (A), between the relative abundance of *S. cerevisiae* and ATEC (B), between the relative abundance of *C. albicans* and ATEC (C), between the relative abundance of *C. albicans* and *S. cerevisiae* (D). p values < 0.05 are considered statistically significant.

## Discussion

The gastrointestinal (GI) tract harbors essential bacteria and fungi that are crucial in maintaining host health ^35^. However, mycobiota, which refers to the fungal component of the microbiota, remains under-explored ^35,36^, especially in the context of neurological disorders like ASD where the gut bacteria have been studied widely ^5^. To fill the gaps for mycobiota in ASD, in this pilot study, we sought to elucidate the composition of fungal communities in children with ASD and analyze the interrelationship between the mycobiota, GI, and ASD symptoms.

Our taxonomic analysis identified a significantly lower abundance of *S. cerevisiae* in children with ASD compared to TD children (Fig. 2A). Multiple reports have shown that *S*. *cerevisiae* and a genetically closely related strain *S. cerevisiae var. boulardii* are prevalent gut commensals and exert salutary effects by acting as probiotics, conferring protection against antibiotic or *C. difficile* induced diarrhea and ameliorating conditions such as gastroenteritis ^37–39^, and ulcerative colitis ^40^ (Sun et al. 2021). Kobliner et al. reported that treatment with *S. cerevisiae var. boulardii* to a child with ASD (a single case report), gastrointestinal dysfunction, obsessive-compulsive disorder (OCD), and self-injurious behavior (SIB), successfully reduced OCD and SIB ^41^. Conversely, Zou et al. reported that *S. cerevisiae* was significantly higher in ASD compared to TD children ^20^.

Interestingly, we observed children with ASD with <22% of the abundance of *S. cerevisiae* had significantly higher autism symptoms (ATEC scores) (Fig. 2D) compared to children with ASD with >22% of the abundance of *S. cerevisiae*, which suggests that *S. cerevisiae* could be beneficial to reduce the autism symptoms ^41^. Also, *S. cerevisiae* was found to anti-correlate with *C albicans*, an opportunistic pathogen (Fig. 4D).

Whereas *S. cerevisiae* was the dominant fungi in TD children, total *Candida* and *C. albicans* were dominant in the gastrointestinal flora of ASD children (Fig. 1A-B). Nonetheless, no statistically significant difference was observed between TD and ASD children for total *Candida* (Table S2) and *C. albicans* (Fig. 2C). *Candida* (yeast) is an opportunistic pathogen and commonly found in the human gut, however, an increased abundance of fungi or yeast (primarily *C. albicans*) has been reported in children with ASD ^3,17–21^. Looking closely at the data, we observed a bimodal distribution of *Candida* and *C. albicans* abundance within ASD children. 17.5% of ASD children with an elevated abundance of total *Candida* (>80%) and 15% of ASD children with a higher abundance of *C. albicans* (>80%) had significantly higher ATEC scores (Fig. 3C-D), which suggests that overgrowth of *Candida* and *C. albicans* and lower abundance of *S. cerevisiae* can worsen autism symptoms (Fig. 2A, D). Previous studies *via* semi-quantitative yeast culture of fecal microbiota that ASD children exhibited higher *Candida* culture positives than TD children ^3,18^. This trend has also been reported for other conditions like irritable bowel syndrome (IBS) ^42^ and Crohn’s Disease ^43^.

Furthermore, one study ^3^ found that the increase in *Candida* in ASD children was associated with diminished levels of secretory Immunoglobulin A (sIgA), which is the primary antibody against bacteria and fungi in the mucous membranes ^44,45^. The depletion of sIgA may be due to the production of IgA-specific proteases secreted by *C. albicans* which degrade sIgA ^46–48^. It has also been postulated that a higher abundance of *Candida* or C. *albicans* can increase the production of ammonia, consequently leading to an excess of gamma-aminobutyric Acid (GABA) production, which might be associated with autism-like behaviors ^49^. *Candida* also produces mycotoxins, such as alcohols, aldehydes, 2-phenylethanol, tartaric acid, isoamyl alcohol, and tryptophol ^23,50–52^, which are known to affect cognitive function adversely ^53^. It is important to note that in some studies, ASD children with a higher abundance of *Candida* suffer from GI issues ^18^. In this study, we observed that GI issues were significantly higher in ASD children than in TD (Fig. S5), and significantly positively correlated with ATEC scores (Fig. 4A) and *C. albicans* (Fig. 4C). However, *C. albicans* and GI symptoms did not show a significant correlation (Fig. S6B). This suggests that elevated levels of yeast/yeast or their metabolites may affect primarily neurological symptoms but not GI symptoms. It has been reported that children with ASD often have increased use of oral antibiotics ^9,15^, which may foster the proliferation of *Candida*. Intriguingly, anti-fungal interventions have been identified as having therapeutic potential in ASD treatment ^32,49,50,54^. Altogether, the above findings suggest that high incidence of yeast infections in children with ASD and an increased abundance of *Candida* or *C*. *albicans* could be involved in the underlying pathophysiology of ASD ^49,54^.

The lack of an association of fungi/yeast with GI symptoms suggests that yeast/fungi dysbiosis can exist undetected, with its only symptom being an exacerbation of ASD symptoms. Yeast/fungi dysbiosis could be self-perpetuating once established, since yeast form colonies and produce proteases that destroy secretory IgA, the primary defense against yeast. Longitudinal studies are needed to examine if yeast/fungi dysbiosis are long-lasting, and if they potentially have a long-lasting effect on ASD symptoms.

A limitation of this study is a small sample size to distinguish the yeast community between ASD and TD cohorts, as well as their interrelation with autism and GI symptoms. This study is based on fungal ITS gene amplicon sequencing, but shotgun metagenomics could shed light on understanding the functional dynamics of the yeast/fungal microbiome in ASD children.

In summary, fungal ITS gene amplicon sequencing revealed that S. *cerevisiae* was significantly lower in ASD vs. TD controls, and anti-correlated with levels of *C. albicans*, a known opportunistic pathogen. Autism severity (ATEC scores) was significantly higher in the ASD group with lower *S. cerevisiae* and higher *C albicans*. However, GI symptoms did not correlate with levels of *S. cerevisiae* or *C albicans*, suggesting that *Candida* or *C*. *albicans* primarily affects neurological/ASD symptoms, but not GI symptoms. Total levels of *Candida* and *C. albicans* were not significantly different in ASD vs. TD controls (unlike previous studies), suggesting that the ASD children are unusually sensitive to the neurological effects of yeast. GI symptoms (6-GSI) were found to positive correlate with autism severity (ATEC), but this correlation appears to be due to non-yeast factors, such as possibly abnormal gut bacteria or GI pain. We hypothesize that early detection of yeast infection/dysbiosis, and treatment with antifungals targeting *Candida* and *C. albicans* can help reduce autism severity, and recommend that an intervention study with antifungals and possibly an anti-yeast diet be conducted. We recommend future research to include a larger cohort, a longitudinal approach, if possible with an intervention, and shotgun metagenomics, to fathom the functional role of fungi/yeast in ASD children.

## Methods

### Enrollment of study participants

A total of 78 participants, encompassing 38 TD children and 40 children with ASD, were enrolled in this study. The age range of the subjects was 3-17 years (Table S1). A critical inclusion criterion was that the subjects did not use any type of antibiotic or antifungal medications for a minimum duration of one month before fecal sample collection. GI symptoms were assessed utilizing a modified 6-item version ^3^ of the original Gastrointestinal Severity Index (GSI) questionnaire with 6 items of GI severity index ^55^. Autism symptoms were assessed with the Autism Treatment Evaluation Checklist (ATEC) to corroborate the diagnosis of ASD. We also evaluated nutritional supplements and special diets, such as gluten-free/casein-free (GF/CF) diet, probiotics use, seafood consumption, and nutrient supplements (e.g., vitamins/calcium) ^8^. For more information, see the supplementary document S1-S2.

### Ethical and institutional Review board information

The study was conducted according to the guidelines of the Declaration of Helsinki and approved by the Institutional Review Board at Arizona State University, AZ, USA (ASU IRB Protocol #: 1206007979, 1004005109). The study was advertised in Arizona, USA. Interested candidates were mailed a consent form (with a preliminary questionnaire and GI severity index scoring) and general participation questions. Upon receiving a signed informed consent form and completed questionnaires from parents/guardians, sample collection kits were mailed to the participants.

### Sample Collection and fecal DNA extraction

Parents/guardians collected fecal samples from their children and immediately froze them. Subsequently, these samples were shipped to Arizona State University with dry ice. Upon receipt at the laboratory, samples were stored at −80°C until DNA extraction. Total genomic DNA was isolated using a PowerSoil DNA extraction kit (Mobio Carlsbad, CA) with a default protocol from the manufacturer.

### Fungal ITS gene amplicon sequencing

The extracted fecal DNA was processed for fungal sequencing using the fungal internal transcribed spacer (ITS) Illumina Amplicon Protocol obtained from the Earth Microbiome Project (http://www.earthmicrobiome.org/protocols-and-standards/its/). The library was constructed using the ITS1f-ITS2 primer set (Forward primer 5’-CTTGGTCATTTAGAGGAAGTAA-3’ and Reverse primer 5’-GCTGCGTTCTTCATCGATGC-3’). Amplicons were subsequently pooled and subjected to purification using the MoBio UltraClean PCR Clean-Up Kit. Finally, an aliquot from the pooled sample was sequenced using an Illumina MiSeq sequencer with single-end forward reads of ~250bp. Sequencing was performed at the Genomics core in the Biodesign Institute of Arizona State University, AZ, USA.

### Sequencing data analyses

Raw sequencing reads were processed using QIIME2 (v2022.02) ^33^. Pre-processing involved removing the PCR primers, DNA barcodes, and the filtration of sequence reads for low-quality nucleotides. Reads were filtered based on phred score >25 and trimmed at 210bp. Denoising was performed using DADA2 ^56^ to remove duplicate reads and chimeric sequences. Amplicon sequence variants (ASV) were assigned using the UNITE database (v8, 10.5.2021) ^34^ The sequences were clustered at a 99% similarity against the UNITE database. Rarefaction was performed at 1,329 reads depth to consider all samples for further downstream analysis. Downstream analyses and visualization of relative abundance and diversity indices were performed using the Dokdo API (https://github.com/sbslee/dokdo, v1.14.0) in Python with Qiime2.

### Quantitative PCR (qPCR) and semi-quantitative fungal abundance

To quantify the total fungal DNA in fecal DNA, a TaqMan qPCR assay was performed using FungiQuant universal primers FungiQuant-F: 5′-GGRAAACTCACCAGGTCCAG-3′ and FungiQuant-R: 5′-GSWCTATCCCCAKCACGA3′, FungiQuant-Prb (6FAM) 5′-TGGTGCATGGCCGTT-3′ (MGBNFQ) ^57^ and QuantStudio3 instrument (Applied Biosystems, CA, USA). We used the following qPCR conditions: 10 min at 95°C for Taq activation, 15 s at 95°C for denaturation, and 1 min at 65°C for annealing and extension x 50 cycles ^57^. All samples and controls (positive and negative) were run in triplicates, and *C. albicans* (ATCC 5314) genomic DNA (0.37 pg/μL) was used as a standard control to generate calibration curves. Total fungal DNA was reported in gene copy number/gm wet feces.

Additionally, semi-quantitative abundance was calculated to identify 18S rRNA genes for specific fungal taxa. This was achieved by multiplying the relative abundance (%) of selected fungal taxa (*C. albicans, S. cerevisiae*) by the resultant copy numbers of the 18S rRNA gene per mg of wet feces in each corresponding sample ^58^. Due to constraints in DNA availability, qPCR was conducted on a subset of the cohort, comprising 28 TD and 20 ASD children.

### Statistical analysis and plots

To investigate and compare the differences in fungal composition (as determined by both sequencing and qPCR), and symptoms between TD and ASD groups, a nonparametric statistical test, the Mann-Whitney U test, was performed, assuming the data were non-normally distributed due to a small sample size. To avoid false positives, all p-values are corrected with the Benjamini-Hochberg method ^59^. Additionally, Pearson and Spearman correlation tests were performed to analyze correlations between taxa and symptoms. Corrected p values <0.05 were considered statistically significant. For the box plot of *C. albicans* and *S. cerevisiae*, a pseudo-count (0.0001) was incorporated for each sample due to zeros (Fig 2A,2C, Fig S4B-C), and subsequently, data was transformed into a log10 scale. All statistical analyses were performed in R (v0.4.0), and figures were generated through R using *ggplot2* (v3.4.2), *ggpubr* (v0.6.0), and OriginPro (v2023b). Inkscape (v1.1) was used to create or edit the figures.

## Acknowledgments

We gratefully thank all children with ASD and their families for participating in the study. We also would like to acknowledge Juan Maldonado and team from the Genomics Core–Biodesign Institute ASU for ITS sequencing and. We thank Mihyun Lee for assisting with the references.

## Authors’ contributions

Conceptualization, K.N., D.-W.K., J.B.A. and R.K.-B.; data curation, K.N., J.P., J.B.A. and R.K.-B.; formal analysis, K.N, J. P., D.-W.K., J.B.A. and R.K.-B.; funding acquisition, J.B.A. and R.K.-B.; investigation, K.N, J. P., D.-W.K., J.B.A. and R.K.-B.; methodology, K.N., J. P., D.-W.K., A. B., J.B.A. and R.K.-B.; project administration, D.-W.K., R.K.-B. and J.B.A.; software, K.N., J., P., and D.-W.K.; supervision, D.-W.K., J.B.A. and R.K.-B.; visualization, K.N. J. P., D.-W.K.; writing—original draft, K.N., J.B.A. and R.K.-B.; writing—review and editing, K.N., J. P., D.-W.K., A. B., D. A. B., J.B.A. and R.K.-B.. All authors have read and agreed to the published version of the manuscript.

## Availability of data and materials

The data presented in this study are openly available in the NCBI SRA repository under BioProject ID PRJNA993699 and can be accessed at https://www.ncbi.nlm.nih.gov/bioproject/993699

## List of abbreviations

ASD: Autism spectrum disorders
ASU: Arizona State University
ATEC: Autism Treatment Effectiveness Checklist
DNA: Deoxyribonucleic acid
CDC: Centers for Disease Control and Prevention
GI: Gastrointestinal
GSI: Gastrointestinal Severity Index
ITS: Internal transcribed spacer
sIgA: secretory Immunoglobulin A
PCR: Polymerase Chain Reaction
QC: Quality Control
qPCR: quantitative Polymerase Chain Reaction
Qiime: Quantitative Insights Into Microbial Ecology
RNA: Ribonucleic acid
TD: Typically-Developing

## References

1. CDC. 2023-ADDM-Factsheet_508. [cited 2023 Jul 8]; Available from: https://www.cdc.gov/ncbddd/autism/materials/addm-factsheet.html

2. Antaki D, Guevara J, Maihofer AX, Klein M, Gujral M, Grove J, Carey CE, Hong O, Arranz MJ, Hervas A, et al. A phenotypic spectrum of autism is attributable to the combined effects of rare variants, polygenic risk and sex. Nat Genet 2022; 54:1284–92.

3. Adams JB, Johansen LJ, Powell LD, Quig D, Rubin RA. Gastrointestinal flora and gastrointestinal status in children with autism - comparisons to typical children and correlation with autism severity. BMC Gastroenterol 2011; 11.

4. Krajmalnik-Brown R, Lozupone C, Kang D-W, Adams JB. Gut bacteria in children with autism spectrum disorders: challenges and promise of studying how a complex community influences a complex disease. Microb Ecol Health Dis 2015; 26.

5. Morton JT, Jin DM, Mills RH, Shao Y, Rahman G, McDonald D, Zhu Q, Balaban M, Jiang Y, Cantrell K, et al. Multi-level analysis of the gut–brain axis shows autism spectrum disorder-associated molecular and microbial profiles. Nat Neurosci 2023;

6. Collins SM, Surette M, Bercik P. The interplay between the intestinal microbiota and the brain. Nat Rev Microbiol 2012; 10:735–42.

7. Mayer EA, Knight R, Mazmanian SK, Cryan JF, Tillisch K. Gut microbes and the brain: Paradigm shift in neuroscience. Journal of Neuroscience 2014; 34:15490–6.

8. Kang DW, Park JG, Ilhan ZE, Wallstrom G, LaBaer J, Adams JB, Krajmalnik-Brown R. Reduced Incidence of Prevotella and Other Fermenters in Intestinal Microflora of Autistic Children. PLoS One 2013; 8.

9. Kang DW, Adams JB, Gregory AC, Borody T, Chittick L, Fasano A, Khoruts A, Geis E, Maldonado J, McDonough-Means S, et al. Microbiota Transfer Therapy alters gut ecosystem and improves gastrointestinal and autism symptoms: An open-label study. Microbiome 2017; 5.

10. Dan Z, Mao X, Liu Q, Guo M, Zhuang Y, Liu Z, Chen K, Chen J, Xu R, Tang J, et al. Altered gut microbial profile is associated with abnormal metabolism activity of Autism Spectrum Disorder. Gut Microbes 2020; 11:1246–67.

11. Finegold SM, Dowd SE, Gontcharova V, Liu C, Henley KE, Wolcott RD, Youn E, Summanen PH, Granpeesheh D, Dixon D, et al. Pyrosequencing study of fecal microflora of autistic and control children. Anaerobe [Internet] 2010 [cited 2022 May 3]; 16:444–53. Available from: https://pubmed.ncbi.nlm.nih.gov/20603222/

12. Iglesias-vázquez L, Riba GVG, Arija V, Canals J. Composition of gut microbiota in children with autism spectrum disorder: A systematic review and meta-analysis. Nutrients 2020; 12.

13. Averina OV., Kovtun AS, Polyakova SI, Savilova AM, Rebrikov DV., Danilenko VN. The bacterial neurometabolic signature of the gut microbiota of young children with autism spectrum disorders. J Med Microbiol 2020; 69:558–71.

14. Ding X, Xu Y, Zhang X, Zhang L, Duan G, Song C, Li Z, Yang Y, Wang Y, Wang X, et al. Gut microbiota changes in patients with autism spectrum disorders. J Psychiatr Res 2020; 129:149–59.

15. Vargason T, McGuinness DL, Hahn J. Gastrointestinal Symptoms and Oral Antibiotic Use in Children with Autism Spectrum Disorder: Retrospective Analysis of a Privately Insured U.S. Population. J Autism Dev Disord 2019; 49:647–59.

16. Gutierrez D, Weinstock A, Antharam VC, Gu H, Jasbi P, Shi X, Dirks B, Krajmalnik-Brown R, Maldonado J, Guinan J, et al. Antibiotic-induced gut metabolome and microbiome alterations increase the susceptibility to Candida albicans colonization in the gastrointestinal tract. FEMS Microbiol Ecol 2020; 96.

17. Kantarcioglu AS, Kiraz N, Aydin A. Microbiota–Gut–Brain Axis: Yeast Species Isolated from Stool Samples of Children with Suspected or Diagnosed Autism Spectrum Disorders and In Vitro Susceptibility Against Nystatin and Fluconazole. Mycopathologia 2016; 181:1–7.

18. Iovene MR, Bombace F, Maresca R, Sapone A, Iardino P, Picardi A, Marotta R, Schiraldi C, Siniscalco D, Serra N, et al. Intestinal Dysbiosis and Yeast Isolation in Stool of Subjects with Autism Spectrum Disorders. Mycopathologia 2017; 182:349–63.

19. Strati F, Cavalieri D, Albanese D, De Felice C, Donati C, Hayek J, Jousson O, Leoncini S, Renzi D, Calabrò A, et al. New evidences on the altered gut microbiota in autism spectrum disorders. Microbiome 2017; 5.

20. Zou R, Wang Y, Duan M, Guo M, Zhang Q, Zheng H. Dysbiosis of Gut Fungal Microbiota in Children with Autism Spectrum Disorders. J Autism Dev Disord 2021; 51:267–75.

21. Alookaran J, Liu Y, Auchtung TA, Tahanan A, Hessabi M, Asgarisabet P, Rahbar MH, Fatheree NY, Pearson DA, Mansour R, et al. Fungi: Friend or Foe? A Mycobiome Evaluation in Children with Autism and Gastrointestinal Symptoms. J Pediatr Gastroenterol Nutr 2022; 74:377–82.

22. Baker S, Shaw W. Case Study: Rapid Complete Recovery From An Autism Spectrum Disorder After Treatment of Aspergillus With The Antifungal Drugs Itraconazole And Sporanox. Integr Med (Encinitas) 2020; 19:20–7.

23. Deitrich RA, Petersen D, Vasiliou V. Removal of acetaldehyde from the body. Novartis Found Symp 2007; 285:23–40; discussion 40-51, 198–9.

24. Ghosh S, Kebaara BW, Atkin AL, Nickerson KW. Regulation of aromatic alcohol production in Candida albicans. Appl Environ Microbiol [Internet] 2008 [cited 2023 Jul 6]; 74:7211–8. Available from: https://pubmed.ncbi.nlm.nih.gov/18836025/

25. Hendler RA, Ramchandani VA, Gilman J, Hommer DW. Stimulant and sedative effects of alcohol. Curr Top Behav Neurosci [Internet] 2013 [cited 2023 Jul 6]; 13:489–509. Available from: https://pubmed.ncbi.nlm.nih.gov/21560041/

26. Karami N, Mirzajani F, Rezadoost H, Karimi A, Fallah F, Ghassempour A, Aliahmadi A. Open Peer Review Discuss this article (0) Comments Initial study of three different pathogenic microorganisms by gas chromatography-mass spectrometry [version 3; referees: 3 approved]. 2018;

27. De Santis B, Brera C, Mezzelani A, Soricelli S, Ciceri F, Moretti G, Debegnach F, Bonaglia MC, Villa L, Molteni M, et al. Role of mycotoxins in the pathobiology of autism: A first evidence. Nutr Neurosci [Internet] 2019 [cited 2023 Jul 6]; 22:132–44. Available from: https://pubmed.ncbi.nlm.nih.gov/28795659/

28. Shaw W, Kassen E, Chaves E. Assessment of Antifungal Drug Therapy in Autism by Measurement of Suspected Microbial Metabolites in Urine with Gas Chromatography-Mass Spectrometry. 2000.

29. Adams JB, Johansen LJ, Powell LD, Quig D, Rubin RA. Gastrointestinal flora and gastrointestinal status in children with autism--comparisons to typical children and correlation with autism severity. BMC Gastroenterol [Internet] 2011 [cited 2023 Jul 6]; 11. Available from: https://pubmed.ncbi.nlm.nih.gov/21410934/

30. Casadevall A, Pirofski LA. Immunoglobulins in defense, pathogenesis, and therapy of fungal diseases. Cell Host Microbe [Internet] 2012 [cited 2023 Jul 6]; 11:447–56. Available from: https://pubmed.ncbi.nlm.nih.gov/22607798/

31. Fadlallah J, Sterlin D, Fieschi C, Parizot C, Dorgham K, El Kafsi H, Autaa G, Ghillani-Dalbin P, Juste C, Lepage P, et al. Synergistic convergence of microbiota-specific systemic IgG and secretory IgA. J Allergy Clin Immunol [Internet] 2019 [cited 2023 Jul 6]; 143:1575–1585.e4. Available from: https://pubmed.ncbi.nlm.nih.gov/30554723/

32. Edelson S, Rimland B. ParentRatings 2009. [cited 2023 Jul 8]; Available from: https://autism.org/treatment-ratings-for-autism/

33. Bolyen E, Rideout JR, Dillon MR, Bokulich NA, Abnet CC, Al-Ghalith GA, Alexander H, Alm EJ, Arumugam M, Asnicar F, et al. Reproducible, interactive, scalable and extensible microbiome data science using QIIME 2. Nat Biotechnol 2019; 37:852–7.

34. Nilsson RH, Larsson KH, Taylor AFS, Bengtsson-Palme J, Jeppesen TS, Schigel D, Kennedy P, Picard K, Glöckner FO, Tedersoo L, et al. The UNITE database for molecular identification of fungi: Handling dark taxa and parallel taxonomic classifications. Nucleic Acids Res 2019; 47:D259–64.

35. Zhang F, Aschenbrenner D, Yoo JY, Zuo T. The gut mycobiome in health, disease, and clinical applications in association with the gut bacterial microbiome assembly. Lancet Microbe 2022; 3:e969–83.

36. Huffnagle GB, Noverr MC. The emerging world of the fungal microbiome. Trends Microbiol 2013; 21:334–41.

37. Rajkowska K, Kunicka-Styczyńska A. Phenotypic and genotypic characterization of probiotic yeasts. Biotechnology and Biotechnological Equipment 2009; 23:662–5.

38. West C, Stanisz AM, Wong A, Kunze WA. Effects of Saccharomyces cerevisiae or boulardii yeasts on acute stress induced intestinal dysmotility. World J Gastroenterol 2016; 22:10532–44.

39. Abid R, Waseem H, Ali J, Ghazanfar S, Ali GM, Elasbali AM, Alharethi SH. Probiotic Yeast Saccharomyces: Back to Nature to Improve Human Health. Journal of Fungi 2022; 8.

40. Sun S, Xu X, Liang L, Wang X, Bai X, Zhu L, He Q, Liang H, Xin X, Wang L, et al. Lactic Acid-Producing Probiotic Saccharomyces cerevisiae Attenuates Ulcerative Colitis via Suppressing Macrophage Pyroptosis and Modulating Gut Microbiota. Front Immunol 2021; 12.

41. Kobliner V, Mumper E, Baker SM. Reduction in Obsessive Compulsive Disorder and Self-Injurious Behavior With Saccharomyces boulardii in a Child with Autism: A Case Report. Integr Med (Encinitas) 2018; 17:38–41.

42. Sokol H, Leducq V, Aschard H, Pham HP, Jegou S, Landman C, Cohen D, Liguori G, Bourrier A, Nion-Larmurier I, et al. Fungal microbiota dysbiosis in IBD. Gut 2017; 66:1039–48.

43. Kumamoto CA. Inflammation and gastrointestinal Candida colonization. Curr Opin Microbiol 2011; 14:386–91.

44. Casadevall A, Pirofski L-A. Immunoglobulins in defense, pathogenesis, and therapy of fungal diseases. Cell Host Microbe 2012; 11:447–56.

45. Fadlallah J, Sterlin D, Fieschi C, Parizot C, Dorgham K, El Kafsi H, Autaa G, Ghillani-Dalbin P, Juste C, Lepage P, et al. Synergistic convergence of microbiota-specific systemic IgG and secretory IgA. J Allergy Clin Immunol 2019; 143:1575–1585.e4.

46. Kett K. [Candida species produce IgA proteases--an important biological property]. Tidsskr Nor Laegeforen 1989; 109:2037.

47. Kapustina OA, Chaĭnikova IN, Kartashova OL. [Resistance of the fungi Candida to human innate immunity factors]. Gig Sanit 2012; :27–9.

48. Doron I, Mesko M, Li X V., Kusakabe T, Leonardi I, Shaw DG, Fiers WD, Lin WY, Bialt-DeCelie M, Román E, et al. Mycobiota-induced IgA antibodies regulate fungal commensalism in the gut and are dysregulated in Crohn’s disease. Nat Microbiol 2021; 6:1493–504.

49. Mehra A, Arora G, Sahni G, Kaur M, Singh H, Singh B, Kaur S. Gut microbiota and Autism Spectrum Disorder: From pathogenesis to potential therapeutic perspectives. J Tradit Complement Med 2022;

50. Shaw W, Kassen E, Chaves E. Assessment of Antifungal Drug Therapy in Autism by Measurement of Suspected Microbial Metabolites in Urine with Gas Chromatography-Mass Spectrometry. 2000.

51. Ghosh S, Kebaara BW, Atkin AL, Nickerson KW. Regulation of aromatic alcohol production in Candida albicans. Appl Environ Microbiol 2008; 74:7211–8.

52. Karami N, Mirzajani F, Rezadoost H, Karimi A, Fallah F, Ghassempour A, Aliahmadi A. Initial study of three different pathogenic microorganisms by gas chromatography-mass spectrometry. F1000Res 2017; 6:1415.

53. De Santis B, Brera C, Mezzelani A, Soricelli S, Ciceri F, Moretti G, Debegnach F, Bonaglia MC, Villa L, Molteni M, et al. Role of mycotoxins in the pathobiology of autism: A first evidence. Nutr Neurosci 2019; 22:132–44.

54. Herman A, Herman AP. Could Candida Overgrowth Be Involved in the Pathophysiology of Autism? J Clin Med 2022; 11.

55. Schneider CK, Melmed RD, Barstow LE, Enriquez FJ, Ranger-Moore J, Ostrem JA. Oral human immunoglobulin for children with autism and gastrointestinal dysfunction: A prospective, open-label study. J Autism Dev Disord 2006; 36:1053–64.

56. Callahan BJ, McMurdie PJ, Rosen MJ, Han AW, Johnson AJA, Holmes SP. DADA2: High-resolution sample inference from Illumina amplicon data. Nat Methods 2016; 13:581–3.

57. Liu CM, Kachur S, Dwan MG, Abraham AG, Aziz M, Hsueh P-R, Huang Y-T, Busch JD, Lamit LJ, Gehring CA, et al. FungiQuant: a broad-coverage fungal quantitative real-time PCR assay. BMC Microbiol 2012; 12:255.

58. Bowes DA, Driver EM, Savic S, Cheng Q, Whisner CM, Krajmalnik-Brown R, Halden RU. Integrated multiomic wastewater-based epidemiology can elucidate population-level dietary behaviour and inform public health nutrition assessments. Nat Food 2023.

59. Benjamini Y, Hochberg Y. Controlling the False Discovery Rate: A Practical and Powerful Approach to Multiple Testing. 1995.

