## Supplementary file S1 for "Bimodal distribution of *Candida albicans* in children with Autism linked with ASD symptoms"

**Supplementary Materials S1**

**Methods**

Participant Recruitment and Demographic Features: A total of seventy-eight (78) participants were enrolled in the study, comprising 38 typically developing (TD) and 40 children with autism spectrum disorders (ASD). The inclusion criteria necessitated participants to be within the age bracket of 2.5 to 17 years, and devoid of exposure to any class of antibiotic or antifungal pharmaceutical agents within the preceding 30 days. TDs who were consanguineous first-degree relatives of individuals diagnosed with ASD were precluded from participation. The Autism Treatment Evaluation Checklist (ATEC) was employed and encompasses four dimensions: 1) verbal communication/language, 2) social interaction aptitude, 3) sensory processing/cognitive acuity, and 4) health and physical comportment. The ATEC composite score is calculated by aggregating the scores across these four domains. Elevated ATEC scores are indicative of the heightened severity of ASD.

Additionally, gastrointestinal symptoms were evaluated employing an adapted version of the Gastro-Intestinal Severity Index (GSI) questionnaire (Schneider et al., 2006). Of the GSI dimensions, six symptom categories were incorporated: constipation, diarrhea, stool consistency, stool odor, flatulence, and abdominal discomfort. A trichotomous rating scale was utilized for each symptom category, and an aggregate score was computed, termed the 6-GI Severity Index (6-GSI). Excluded dimensions encompassed ‘unaccounted daytime irritability’, ‘nocturnal arousal’, and ‘abdominal sensitivity’.

### Table S1:

**Table S1.** Characteristics of study participants

| Category | TD | ASD |
|---|---|---|
| Total Subjects (n) | 38 | 40 |
| Male/Female | 29 / 9 | 37 / 3 |
| Age (years) | 8.24 ± 3.78 | 8.65 ± 3.93 |
| ATEC | N/A | 67.82 ± 26.79 |
| PDD-BI | N/A | -55.13 ± 60.24 |
| 6-GSI | 1.11 ± 1.69 | 4.7 ± 2.61 |

Data are shown with mean ± standard deviation. TD - Typically Developing, ASD - Autism Spectrum Disorder, ATEC - Autism Treatment Evaluation Checklist, PDD-BI - Pervasive Developmental Disorder Behavior Inventory, GSI - Gastrointestinal Severity Index, N/A - Not Applicable.

**Table S2: Univariate comparison of fungal taxa between TD and ASD children.**

| Name | TD | ASD | p-values |
|---|---|---|---|
| Total *Candida* | 8.77E-02 ± 2.22E-01 | 2.13E-01 ± 3.83E-01 | 0.83 |
| Non-Candida_Total_fungi | 9.12E-01 ± 2.22E-01 | 7.87E-01 ± 3.83E-01 | 0.83 |
| Non-*C.albicans*_Total_fungi | 9.36E-01 ± 2.09E-01 | 8.37E-01 ± 3.59E-01 | 0.67 |
| Non-*C.albicans*_Total_*Candida* | 2.32E-02 ± 9.28E-02 | 5.04E-02 ± 1.84E-01 | 0.64 |
| *s_Saccharomyces_cerevisiae* | 5.19E-01 ± 4.19E-01 | 2.91E-01 ± 3.69E-01 | 0.011 |
| *s_Candida_albicans* | 6.45E-02 ± 2.09E-01 | 1.63E-01 ± 3.59E-01 | 0.67 |
| *g_Penicillium* | 3.75E-02 ± 1.68E-01 | 1.79E-03 ± 4.13E-03 | 0.48 |
| *s_Candida_sake* | 1.88E-02 ± 9.08E-02 | 2.40E-02 ± 1.52E-01 | 0.29 |
| *g_Cladosporium* | 5.05E-03 ± 1.61E-02 | 3.59E-02 ± 1.59E-01 | 0.34 |
| *s_Parastagonospora_nodorum* | 4.72E-05 ± 1.41E-04 | 3.93E-02 ± 1.72E-01 | 0.25 |
| *s_Cyberlindnera_jadinii* | 8.39E-03 ± 4.50E-02 | 6.71E-02 ± 2.22E-01 | 0.71 |
| *s_Penicillium_carneum* | 1.21E-02 ± 7.08E-02 | 1.87E-02 ± 1.18E-01 | 0.60 |
| *s_Candida_tropicalis* | 4.37E-03 ± 2.28E-02 | 2.64E-02 ± 1.09E-01 | 0.51 |
| *s_Aspergillus_heterocaryoticus* | 1.56E-02 ± 5.19E-02 | 1.62E-02 ± 9.78E-02 | 0.052 |
| *s_Issatchenkia_orientalis* | 2.44E-02 ± 1.04E-01 | 3.58E-03 ± 2.04E-02 | 0.40 |
| *g_Wallemia* | 2.25E-02 ± 1.37E-01 | 6.94E-03 ± 3.93E-02 | 0.74 |
| *s_Dekkera_bruxellensis* | 1.70E-03 ± 4.18E-03 | 2.82E-02 ± 1.49E-01 | 0.77 |
| *o_Saccharomycetales* | 2.17E-02 ± 8.84E-02 | 9.00E-03 ± 4.92E-02 | 0.64 |
| *f_Sclerotiniaceae* | 1.09E-02 ± 6.16E-02 | 3.66E-04 ± 2.16E-03 | 0.66 |
| *s_Alternaria_angustiovoidea* | 2.29E-02 ± 1.33E-01 | 1.60E-02 ± 5.05E-02 | 0.26 |
| *s_Candida_vartiovaarae* | 0.00E+00 ± 0.00E+00 | 1.70E-05 ± 1.08E-04 | 0.34 |
| *s_Candida_ethanolica* | 1.66E-05 ± 8.30E-05 | 0.00E+00 ± 0.00E+00 | 0.14 |

In this table, the Mann-Whitney U test was used to compare fungal taxa between TD and ASD children. Data are shown as mean±standard deviation. $p<0.05$ was considered statistically significant. TD - typically developing, ASD - autism spectrum disorder, s - species, g - genera, o - order, f -family.

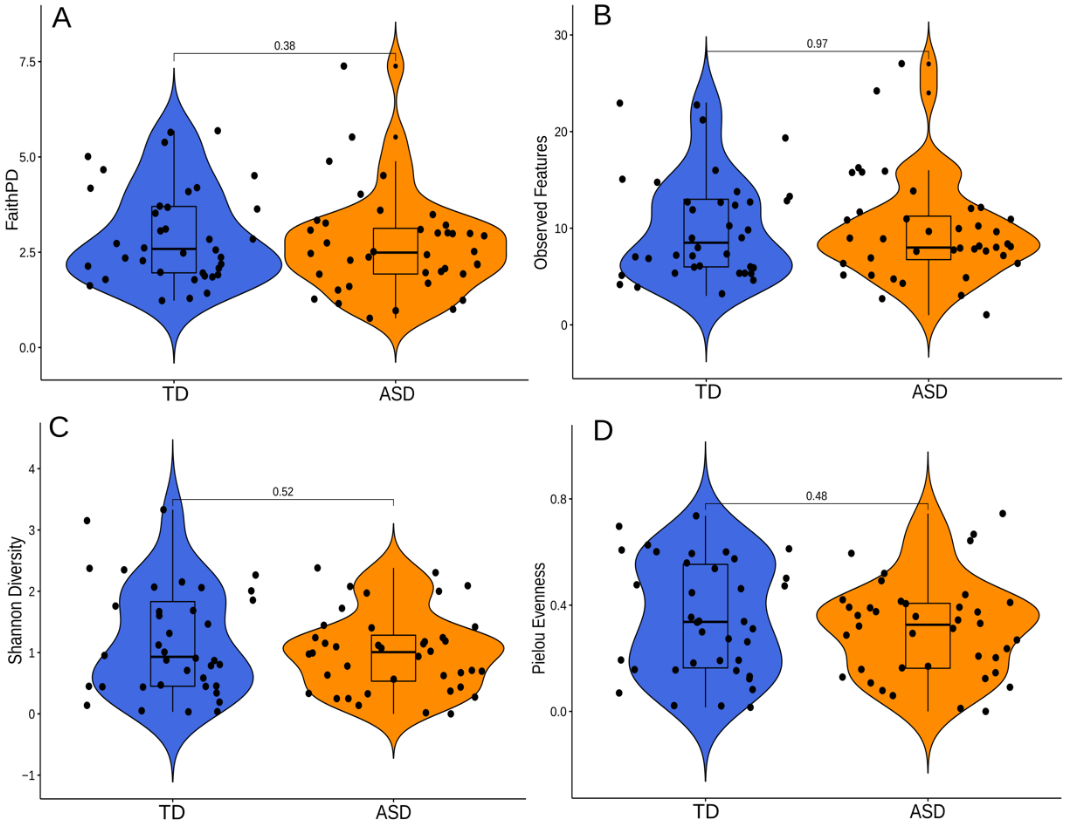


Figure S1: The alpha-diversity indices Faith PD (A), observed features (B), Shannon diversity (C) and Pielou evenness (D) comparisons between TD and ASD children. Blue color represents TD and orange color for ASD participants. p values < 0.05 are considered statistically significant. ASD: autism spectrum disorder; TD: typically developing.

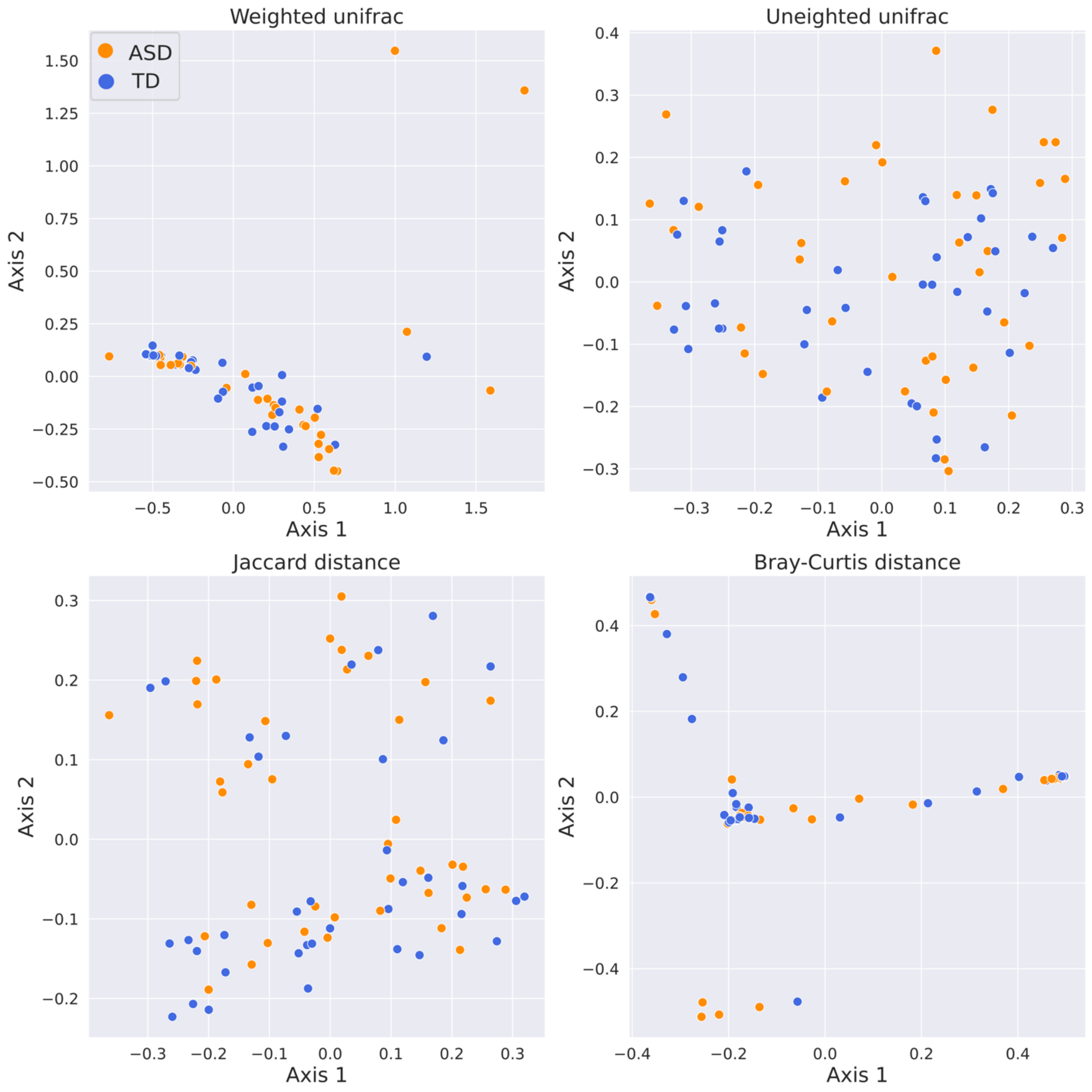


Figure S2: The 2D beta-diversity indices Weighted Unifrac (A), Unweighted Unifrac (B), Jaccard distance (C) and Bray-Curtis distance (D) comparisons between TD and ASD children. Blue color represents TD and orange color for ASD participants. ASD: autism spectrum disorder; TD: typically developing.

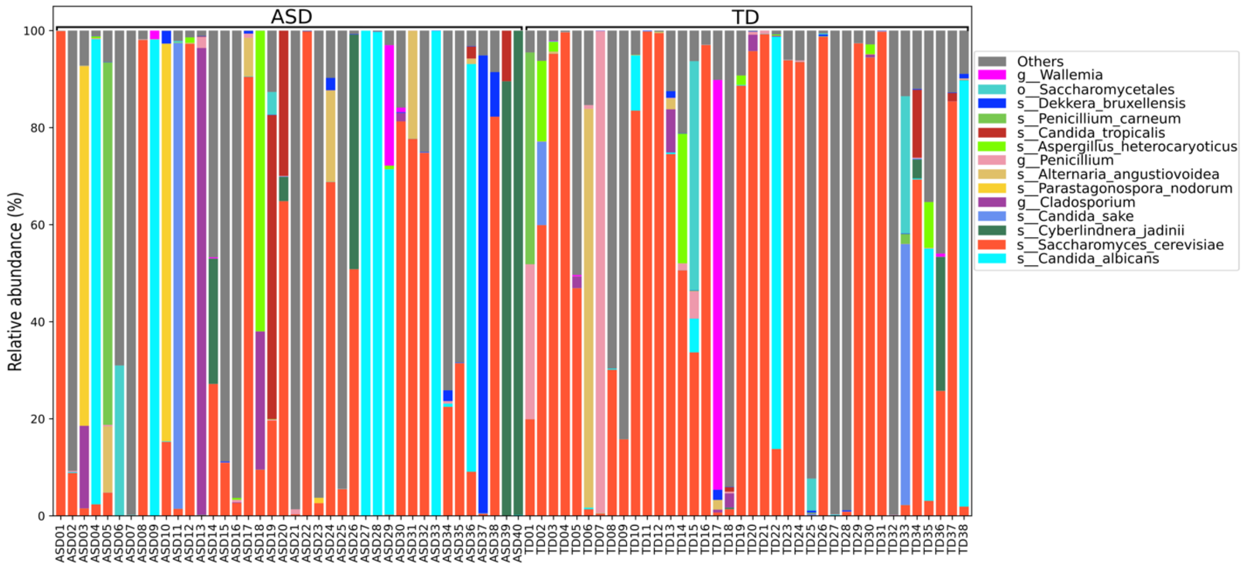


Figure S3: Average relative abundance (%) of Top 15 fungal taxa in TD and children with ASD. TD- typically developing; ASD- autism spectrum disorder. Average relative abundance of fungi in each participant, refer to Figure 1.

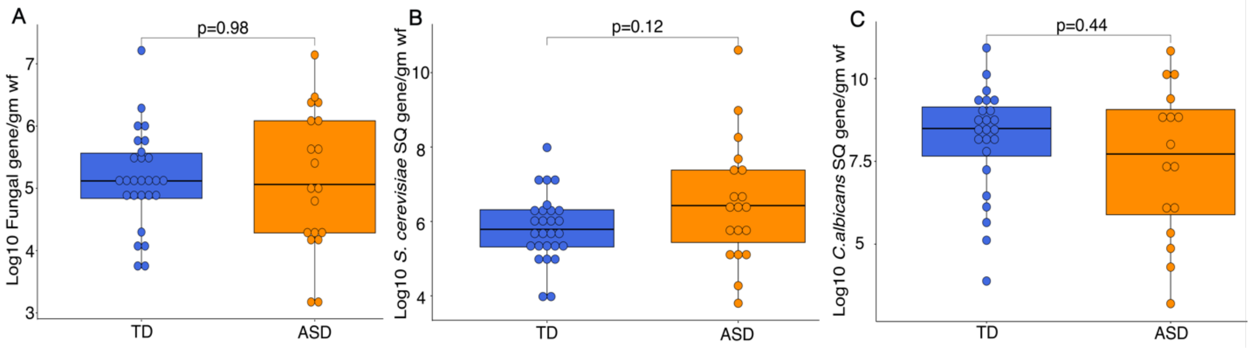


Figure S4: Quantification of the fecal fungal gene in TD and ASD children. Quantitative measurement of total fungal18S rRNA gene (log10) copy/gm wet feces (A), semi-quantitative gene abundance/gm wet feces of S. cerevisiae (B), and C. albicans (C). Blue color represents TD, and orange color for ASD participants. p values < 0.05 are considered statistically significant. ASD: autism spectrum disorder; TD: typically developing, SQ: semi-quantitative, wf: wet feces.

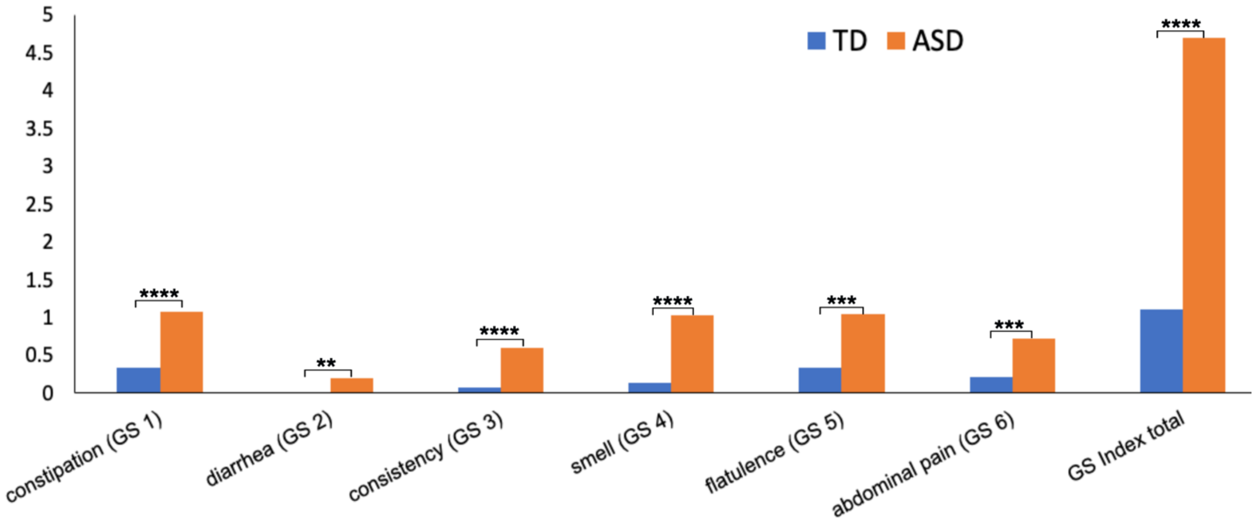


Figure S5: Univariate comparison of gastrointestinal (GI) sub-scores and total between TD and ASD children. Blue color represents TD and orange color for ASD participants.

* Single asterisk indicates $p < 0.05$, ** double asterisks indicate $p < 0.01$, triple *** asterisks indicate $p < 0.001$ and four **** indicate $p < 0.0001$. ASD: autism spectrum disorder; TD: typically developing.

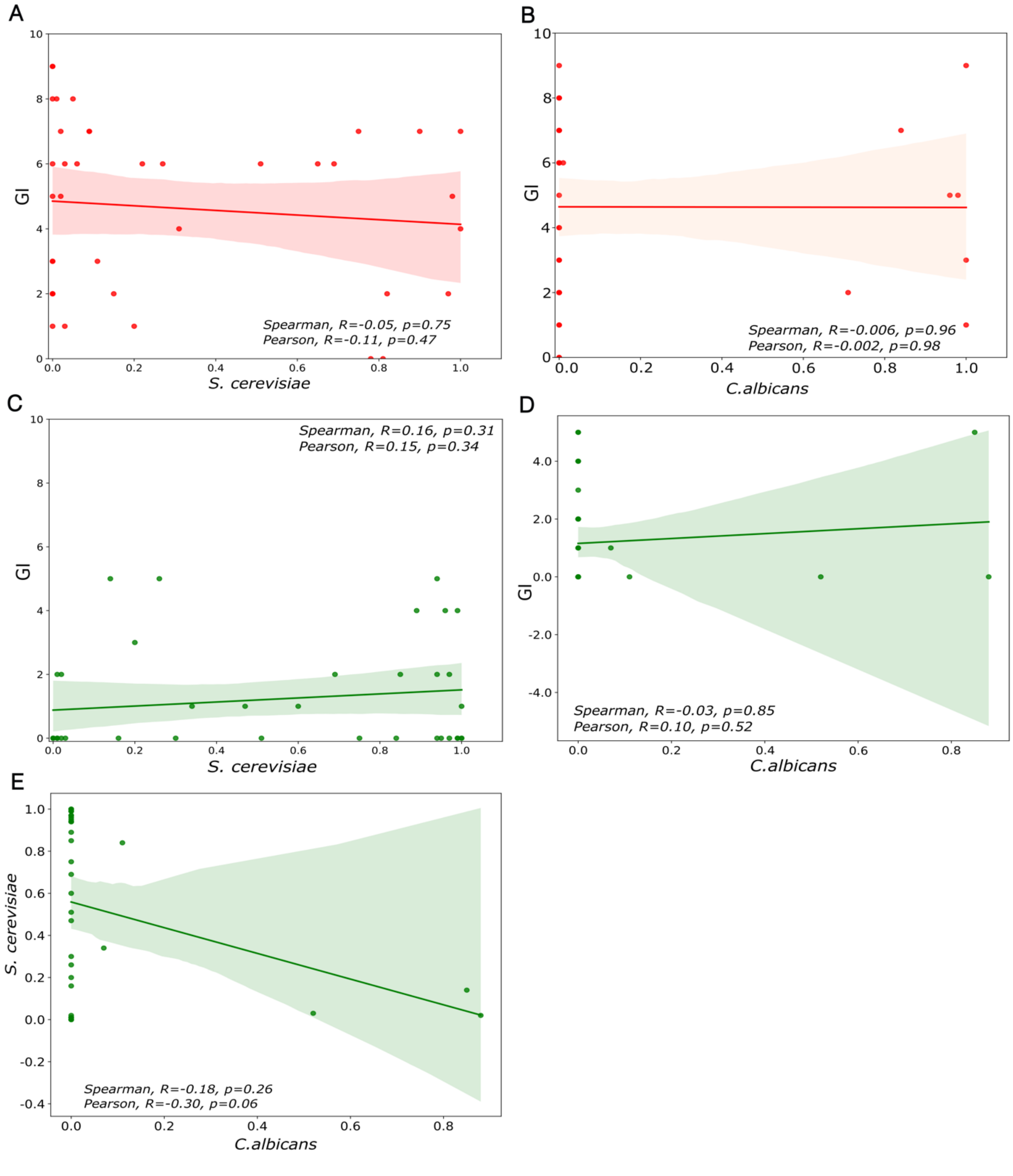


Figure S6: Correlation analyses between fungal taxa and GI symptoms in TD and ASD children. Correlation between GI scores and *S. cerevisiae* (A), and between GI scores and *C. albicans* in ASD children (B), between GI scores and *S. cerevisiae* (C), between GI scores and *C. albicans* (D), and between *C. albicans* and *S. cerevisiae* in TD children (D). p values < 0.05 are considered statistically significant.
